# Whole-brain dynamics and hormonal shifts throughout women’s lifespan: From reproductive stages to menopausal transition and beyond

**DOI:** 10.1101/2024.09.23.614341

**Authors:** Anira Escrichs, Daniela Avila-Varela, Gustavo Patow, Belinda Pletzer, Petra Ritter

## Abstract

Neuroimaging studies have identified significant age-related disruptions in whole-brain dynamics, yet the influence of women’s reproductive stages and associated hormonal shifts remains underexplored. This study leverages resting-state fMRI data from the Human Connectome Project in Aging to examine brain dynamics through five reproductive stages: reproductive, late reproductive, perimenopause, early postmenopause, and late postmenopause. Our results indicate that the late reproductive stage is characterized by the highest dynamical complexity across whole-brain and resting-state networks, while brain dynamics significantly decline at menopause onset. Additionally, we employ machine learning classifiers using two approaches: (1) brain dynamics alone and (2) brain dynamics combined with follicle-stimulating hormone (FSH) and estradiol (brain-hormone model). Both models accurately distinguished reproductive stages, but the brain-hormone model outperformed the brain dynamics model. Key predictors included decreased estradiol, increased FSH, and altered brain dynamics in later life stages. These results offer a framework for assessing brain health across women’s reproductive lifespan.

## Introduction

By 2030, the population of menopausal and postmenopausal women worldwide is expected to reach 1.2 billion, with an annual increase of 47 million women^1^. The menopause transition, also known as perimenopause, is a neurological transition state characterized by declining levels of estrogen and progesterone^2^. While the average age of menopause is typically between 45-55 years, perimenopause can begin 8-10 years before menopause. The neurological symptoms associated with perimenopause, such as hot flashes, irregular menstrual cycles, memory deficits, depression, and insomnia, are caused by disruptions in estrogen-regulated systems that modulate brain function^2^. During perimenopause, the uncoupling of the estrogen receptor network from the bioenergetic system increases the risk of neurodegenerative diseases, particularly Alzheimer’s disease^3–8^. The menopause transition is considered complete when 12 months have passed since the last menstruation, marking postmenopause and reproductive senescence. The age at which a woman experiences her final menstrual period is of significant clinical and public health interest, as it may serve as a potential marker of aging and health^9^.

Recent neuroimaging studies have revealed significant age-related changes in brain dynamics^10–13^. However, these investigations have largely overlooked the influence of women’s reproductive stages, which may significantly affect brain dynamics. Neuroimaging research in women’s health remains considerably limited^14^, and the adaptive responses of the brain to hormonal shifts during peri- and post-menopause remain largely unexplored^15^. Only a few functional MRI studies have focused on peri- and post-menopausal women, typically employing average (static) functional connectivity approaches, seed-based analyses, and small sample sizes^4,16–21^. While previous investigations have provided useful insights, studying the underlying whole-brain dynamics in large neuroimaging datasets can enhance our understanding of how brain networks reconfigure across women’s lifespans.

The brain is inherently a dynamic organ, and analyzing its activity patterns and connectivity over time is required for comprehending how different brain areas interact and communicate to process information efficiently. Static approaches miss these temporal patterns, whereas dynamic analysis can capture brain activity at multiple spatiotemporal scales, offering a more complete understanding of brain function^22–24^. Integrating brain dynamics with female sex hormone levels in machine learning models can enhance our understanding of the brain-hormone interactions that occur throughout women’s reproductive stages and their transitions.

This study used the Lifespan Human Connectome Project in Aging (HCP-A) neuroimaging dataset to explore whole-brain dynamics throughout women’s reproductive stages. Women were classified into five reproductive stages, i.e., reproductive, late reproductive, perimenopause, early postmenopause, and late postmenopause, according to the Reproductive Aging Workshop (STRAW+10) working group guidelines^25^. We hypothesized that (1) significant changes in brain dynamics occur throughout women’s lifespans in relation to their reproductive stage, and (2) machine learning algorithms can accurately classify women into these reproductive stages using a combination of brain dynamics metrics and hormonal biomarkers.

## Results

### Participants

We analyzed resting-state fMRI data of 331 women from the HCP Aging (HCP-A) dataset^26^ (see Methods for a detailed sample description). We obtained the reproductive stage from the data structure called “mchq01”. The HCP-A classifies reproductive stages using the Menstrual Questionnaire, Menopause Screener, serum estradiol and follicle-stimulating hormone (FSH), following the STRAW+10 model ^25^. This model does not use chronological age to determine reproductive staging. **Table 1** shows the characteristics of the sample. Note that some women had fMRI data but not hormone information.

**Table 1.** Sample description. Sample size, average age in years, and standard deviation for each reproductive stage.

|  | **Reproductive** | **Late**<br>**Reproductive** | **Perimenopause** | **Early**<br>**Postmenopause** | **Late**<br>**Postmenopause** |
| --- | --- | --- | --- | --- | --- |
| **Overall sample** |  |  |  |  |  |
| Sample size, N=331 | 56 | 27 | 46 | 35 | 167 |
| Age mean (SD) | 42.51 (4.51) | 43.07 (4.11) | 48.38 (5.45) | 54.10 (5.68) | 71.74 (11.56) |
| Age range | 36.17-55.25 | 36.25-50.92 | 37.42-60.92 | 37.50-61.58 | 40.75-100 |
| **Sample with hormones** |  |  |  |  |  |
| Sample size, N=287 | 48 | 26 | 37 | 34 | 142 |
| Age mean (SD) | 42.32 (4.67) | 43.28 (4.04) | 48.53 (5.31) | 54.11 (5.77) | 72.14 (11.23) |
| Estradiol mean (SD) | 107.79 (110.61) | 113.48 (119.07) | 99.97 (108.18) | 22.28 (31.31) | 9.68 (16.64) |
| FSH mean (SD) | 10.11 (11.78) | 9.40 (8.27) | 32.03 (34.25) | 67.49 (33.60) | 72.39 (27.63) |

### Hormonal biomarkers

The analysis of hormonal biomarkers across reproductive stages revealed distinct patterns **Figure 1A**. FSH levels showed a clear upward trend, with a marked increase from perimenopause to early postmenopause. Conversely, estradiol levels displayed an inverse relationship, declining substantially during perimenopause and reaching their lowest levels in late postmenopause. These profiles align with established endocrine changes in reproductive aging^25^.

**Figure 1.**
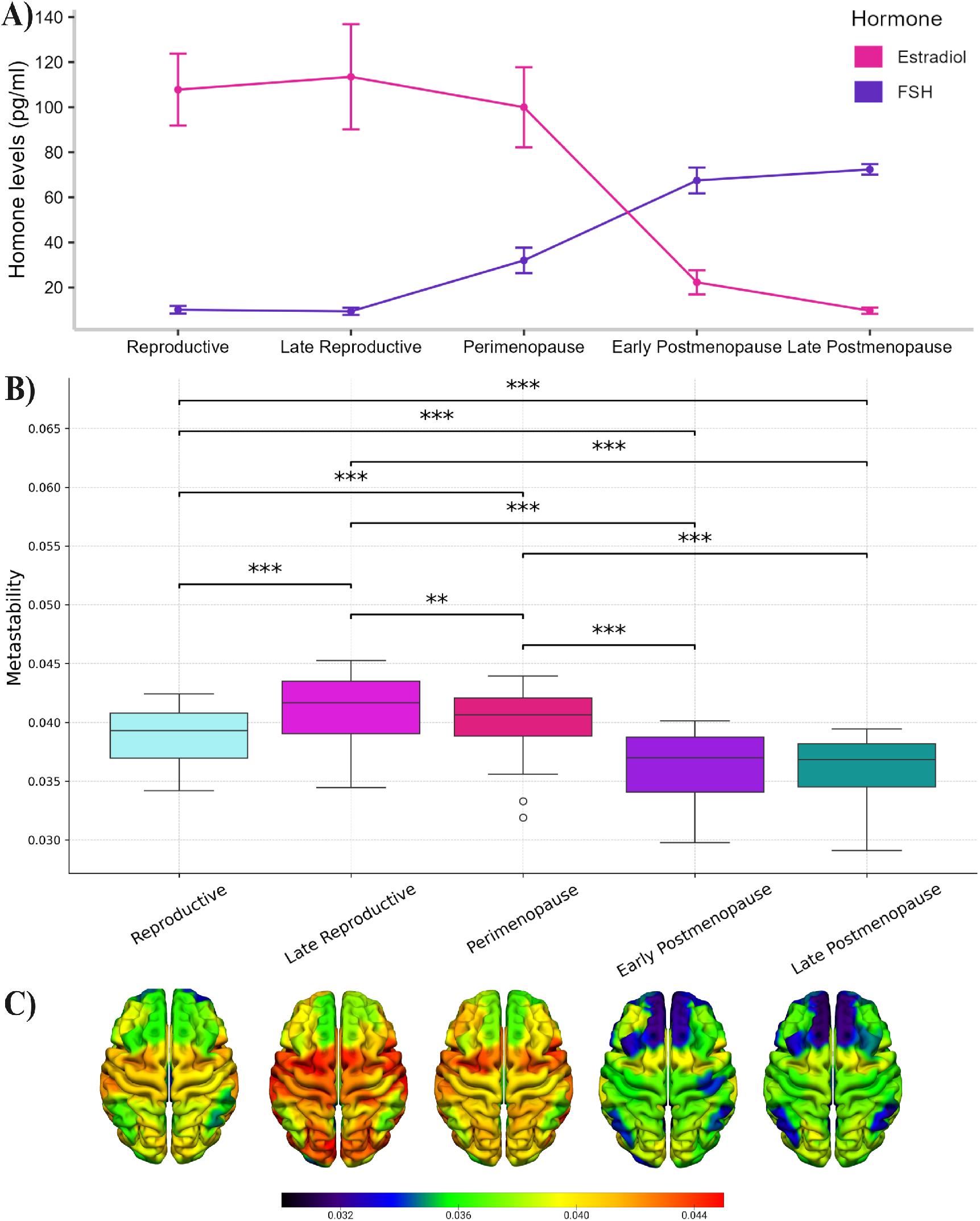
Hormonal profiles and brain dynamics across reproductive stages. A) Mean values (points) and standard errors (whiskers) of FSH and estradiol levels among participants, categorized into five reproductive stages according to STRAW+10 criteria^25^. Estradiol levels decrease and FSH levels increase as women transition from the late reproductive stage to early post-menopause. B) Dynamical complexity across reproductive stages. The late reproductive stage shows the highest dynamical complexity compared to the reproductive, perimenopausal, early postmenopausal, and late postmenopausal stages. Women in the reproductive stage exhibit greater dynamical complexity than those in the early and late postmenopausal stages but less than those in the perimenopausal stage. Additionally, perimenopausal women display higher dynamical complexity compared to women in the early and late postmenopausal stages. The most significant decline in whole-brain dynamics occurs during the critical transition from perimenopause to the onset of menopause, independent of age. This shift aligns with decreasing estradiol and increasing FSH levels (panel A). C) Cortical representations of dynamical complexity for each reproductive stage. Detailed significance levels: *** p < 0.001, ** p < 0.01.

### Dynamical complexity across the whole-brain network

We assessed the dynamical complexity of the whole-brain functional network at each reproductive stage. Specifically, we employed a measure based on metastability to evaluate how spontaneous neural activity in a given node influences the activity of the network over time (more details in methods). Metastability reflects the variability and diversity within a node. Higher metastability indicates increased variability and diversity over time and, thus, increased dynamical complexity. In particular, for each node of the brain parcellation, we computed the average node-metastability over all women in that reproductive stage group. The results depicted in **Figure 1B-C** show that the late reproductive stage displayed the highest dynamical complexity compared to the reproductive (p < 0.001, Monte Carlo permutation and FDR-corrected), perimenopause (p < 0.01, Monte Carlo permutation and FDR-corrected), early postmenopause (p < 0.001, Monte Carlo permutation and FDR-corrected), and late postmenopause (p < 0.001, Monte Carlo permutation and FDR-corrected). Women in the reproductive stage showed higher dynamical complexity than women classified in the early and late postmenopause stages (p < 0.001, Monte Carlo permutation and FDR-corrected) but lower than perimenopausal women (p < 0.001, Monte Carlo permutation and FDR-corrected). Women in perimenopause showed increased dynamical complexity than early and late postmenopausal women (p < 0.001, Monte Carlo permutation and FDR-corrected). These findings reveal a unique pattern of dynamical complexity throughout different stages of reproductive aging, indicating a clear link between female sex hormones and whole-brain network dynamics.

### Dynamical complexity across resting-state networks

We computed the dynamical complexity within 7 well-known resting-state networks^27^ (visual, somatomotor, dorsal attention, salience, limbic, frontoparietal control, and DMN) for each woman in each reproductive stage **(Figure 2)**.

**Figure 2.**
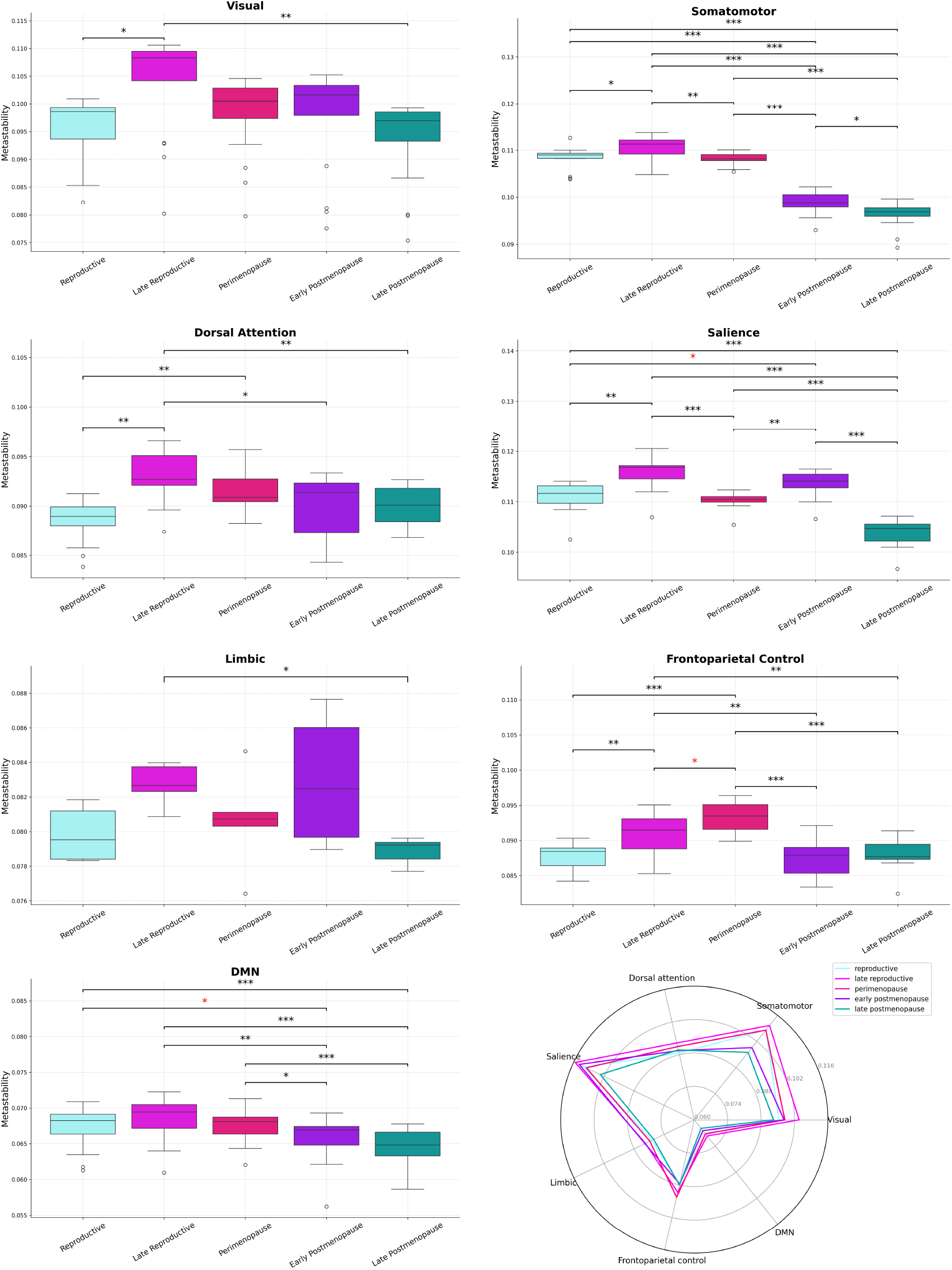
Dynamical complexity within seven resting-state networks across reproductive stages. The late reproductive stage generally showed the highest dynamics across most networks. Interestingly, perimenopausal women exhibited increased dynamical complexity in the control network compared to all other stages. The box in the boxplots indicates the upper and lower quartiles and the line inside indicates the median. The radar plot at the bottom right represents the average metastability values per resting-state network for each reproductive stage. *** represents p < 0.001, ** p < 0.01, * p < 0.05, and * in red marginally significant.

In the visual network, women in the late reproductive state showed higher dynamics compared to the reproductive (p < 0.5, Monte Carlo permutation and FDR-corrected) and late postmenopause (p < 0.01, Monte Carlo permutation and FDR-corrected).

In the somatomotor network, women in the late reproductive state exhibited significantly higher dynamics compared to the reproductive (p < 0.05, Monte Carlo permutation and FDR-corrected), perimenopause (p < 0.01, Monte Carlo permutation and FDR-corrected), early postmenopause (p < 0.001, Monte Carlo permutation and FDR-corrected), and late postmenopause (p < 0.001, Monte Carlo permutation and FDR-corrected). Women in the reproductive stage exhibited greater dynamics than those in the early and late postmenopause stages (p < 0.001, Monte Carlo permutation and FDR-corrected). Perimenopausal women displayed increased dynamical complexity compared to early and late postmenopausal women (p < 0.001, Monte Carlo permutation and FDR-corrected). Women in early postmenopause showed increased dynamics than late postmenopausal women (p < 0.05, Monte Carlo permutation and FDR-corrected).

In the dorsal attention network, women in the late reproductive stage had significantly higher dynamics compared to the reproductive (p < 0.01, Monte Carlo permutation and FDR-corrected), early postmenopause (p < 0.05, Monte Carlo permutation and FDR-corrected), and late postmenopause (p < 0.01, Monte Carlo permutation and FDR-corrected). Women in the reproductive stage exhibited lower dynamics than those in perimenopause (p < 0.01, Monte Carlo permutation and FDR-corrected).

For the salience attention network, women in the late reproductive stage exhibited higher dynamics compared to the reproductive (p< 0.01, Monte Carlo permutation and FDR-corrected), perimenopause (p < 0.001, Monte Carlo permutation and FDR-corrected), and late postmenopause (p < 0.001, Monte Carlo permutation and FDR-corrected). Women in the reproductive stage exhibited lower dynamics than those in early postmenopause (p = 0.056, marginally significant after Monte Carlo permutation and FDR correction) and higher than late postmenopause (p < 0.001, Monte Carlo permutation and FDR-corrected). Perimenopausal women displayed decreased complexity than early postmenopause (p < 0.01, Monte Carlo permutation and FDR-corrected) and increased compared to late postmenopausal women (p < 0.001, Monte Carlo permutation and FDR-corrected). Women in early postmenopause showed greater dynamics than late postmenopause (p < 0.001, Monte Carlo permutation and FDR-corrected).

For the limbic network, women in the late reproductive stage exhibited higher dynamics compared to women in late postmenopause (p < 0.05, Monte Carlo permutation and FDR-corrected).

Interestingly, in the frontoparietal control network, women in perimenopause showed increased dynamics compared to the other groups: reproductive (p < 0.001, Monte Carlo permutation and FDR-corrected), late reproductive (p = 0.08, marginally significant after Monte Carlo permutation and FDR correction), early postmenopause (p < 0.001, Monte Carlo permutation and FDR-corrected), and late postmenopause (p < 0.001, Monte Carlo permutation and FDR-corrected). Women in the late reproductive stage exhibited higher dynamics compared to reproductive (p < 0.01, Monte Carlo permutation and FDR-corrected), early postmenopause (p < 0.01, Monte Carlo permutation and FDR-corrected), and late postmenopause stages (p < 0.01, Monte Carlo permutation and FDR-corrected).

Finally, for the DMN, women in the late reproductive stage showed increased dynamics compared to early postmenopause (p < 0.01, Monte Carlo permutation and FDR-corrected) and late postmenopause (p < 0.001, Monte Carlo permutation and FDR-corrected). Women in the reproductive stage presented increased dynamics with respect to women in the early postmenopause (p = 0.067, marginally significant after Monte Carlo permutation and FDR correction) and late postmenopause stages (p < 0.001, Monte Carlo permutation and FDR-corrected, respectively). Perimenopausal women showed higher dynamical complexity than early and late postmenopausal women (p < 0.05 and p < 0.001, Monte Carlo permutation and FDR-corrected, respectively).

These results indicate that women in the late reproductive stage generally exhibit the highest dynamical complexity, followed by those in the reproductive and perimenopausal stages. This trend shows the significant influence of reproductive aging on brain network dynamics, with notable decreases in dynamical complexity observed in the early and late postmenopausal stages. These findings indicate that reproductive aging is closely linked to brain dynamic reconfigurations across resting-state networks.

### Machine learning classifiers

We tested two approaches. The first approach (brain dynamics model) includes 100 nodes for each woman, and 2) the second approach includes FSH, estradiol and brain dynamics (brain-hormone model). For each approach, we employed Random Forest (RF) and XGBoost (XGB) classifiers^28,29^ to differentiate between reproductive stages (more details in Methods). **Table 2** and **Table 3** summarize the performance metrics, including classifier, accuracy, precision, F1 score, and AUC.

**Table 2.** Performance of machine learning classifiers across groups (brain dynamics model). Note: RF = Random Forest, XGB = XGBoost.

| Hormone Groups | Classifier | Accuracy | Precision | F1 Score | AUC |
| --- | --- | --- | --- | --- | --- |
| Repro. vs Late Repro. | RF | 0.7332 | 0.7333 | 0.7318 | 0.8086 |
|  | XGB | 0.7866 | 0.7890 | 0.7851 | 0.8622 |
| Repro. vs Perimeno | RF | 0.5992 | 0.5985 | 0.5979 | 0.6020 |
|  | XGB | 0.5553 | 0.5538 | 0.5530 | 0.5430 |
| Repro. vs Early Postmeno | RF | 0.7229 | 0.7250 | 0.7227 | 0.7978 |
|  | XGB | 0.7854 | 0.7857 | 0.7857 | 0.8287 |
| Repro. vs Late Postmeno | RF | 0.9013 | 0.9016 | 0.9012 | 0.9433 |
|  | XGB | 0.8682 | 0.8702 | 0.8681 | 0.9226 |
| Late Repro. vs Perimeno | RF | 0.7082 | 0.7066 | 0.7065 | 0.7310 |
|  | XGB | 0.6965 | 0.6960 | 0.6955 | 0.7216 |
| Late Repro. vs Early Postmeno | RF | 0.7571 | 0.7574 | 0.7571 | 0.7261 |
|  | XGB | 0.7000 | 0.7002 | 0.6999 | 0.6971 |
| Late Repro. vs Late Postmeno | RF | 0.9431 | 0.9435 | 0.9431 | 0.9813 |
|  | XGB | 0.9402 | 0.9417 | 0.9401 | 0.9861 |
| Perimeno vs Early Postmeno | RF | 0.7263 | 0.7284 | 0.7282 | 0.7379 |
|  | XGB | 0.6509 | 0.6525 | 0.6520 | 0.6557 |
| Perimeno vs Late Postmeno | RF | 0.8893 | 0.8893 | 0.8892 | 0.9478 |
|  | XGB | 0.8892 | 0.8932 | 0.8889 | 0.9487 |
| Early Postmeno vs Late Postmeno | RF | 0.9313 | 0.9313 | 0.9311 | 0.9779 |
|  | XGB | 0.9071 | 0.9137 | 0.9068 | 0.9689 |

**Table 3.** Performance of machine learning classifiers across groups (brain-hormone model). Note: RF = Random Forest, XGB = XGBoost.

| Hormone Groups | Classifier | Accuracy | Precision | F1 Score | AUC |
| --- | --- | --- | --- | --- | --- |
| Repro. vs Late Repro. | RF | 0.6530 | 0.6558 | 0.6495 | 0.7468 |
|  | XGB | 0.7237 | 0.7239 | 0.7230 | 0.7433 |
| Repro. vs Perimeno | RF | 0.6257 | 0.6252 | 0.6249 | 0.6164 |
|  | XGB | 0.6700 | 0.6723 | 0.6683 | 0.6502 |
| Repro. vs Early Postmeno | RF | 0.9016 | 0.9029 | 0.9017 | 0.9082 |
|  | XGB | 0.8921 | 0.8949 | 0.8927 | 0.9048 |
| Repro. vs Late Postmeno | RF | 0.9402 | 0.9402 | 0.9401 | 0.9875 |
|  | XGB | 0.9462 | 0.9462 | 0.9461 | 0.9739 |
| Late Repro. vs Perimeno | RF | 0.6772 | 0.6742 | 0.6738 | 0.6399 |
|  | XGB | 0.7310 | 0.7284 | 0.7282 | 0.7453 |
| Late Repro. vs Early Postmeno | RF | 0.8429 | 0.8431 | 0.8428 | 0.8498 |
|  | XGB | 0.8571 | 0.8619 | 0.8567 | 0.8918 |
| Late Repro. vs Late Postmeno | RF | 0.9850 | 0.9852 | 0.9850 | 0.9976 |
|  | XGB | 0.9760 | 0.9763 | 0.9760 | 0.9943 |
| Perimeno vs Early Postmeno | RF | 0.7053 | 0.7074 | 0.7062 | 0.7587 |
|  | XGB | 0.7392 | 0.7410 | 0.7386 | 0.7576 |
| Perimeno vs Late Postmeno | RF | 0.9042 | 0.9051 | 0.9041 | 0.9709 |
|  | XGB | 0.9193 | 0.9210 | 0.9191 | 0.9737 |
| Early Postmeno vs Late Postmeno | RF | 0.9223 | 0.9227 | 0.9221 | 0.9681 |
|  | XGB | 0.8953 | 0.8984 | 0.8950 | 0.9603 |

Overall, the brain-hormone model demonstrates stronger classification across most reproductive groups, with higher performance metrics, including accuracy, precision, F1 score, and AUC. The brain-hormone model outperformed the brain dynamics model in 7 out of 10 comparisons. This indicates that incorporating hormonal data alongside brain dynamics enhances the ability of the model to distinguish between different reproductive stages.

### Feature importance

We extracted the top 20 most important features from the brain-hormone model for each pairwise comparison **(Figure 3)**. For clarity, we only present the comparisons between consecutive stages across the transition from reproductive to late postmenopause.

**Figure 3.**
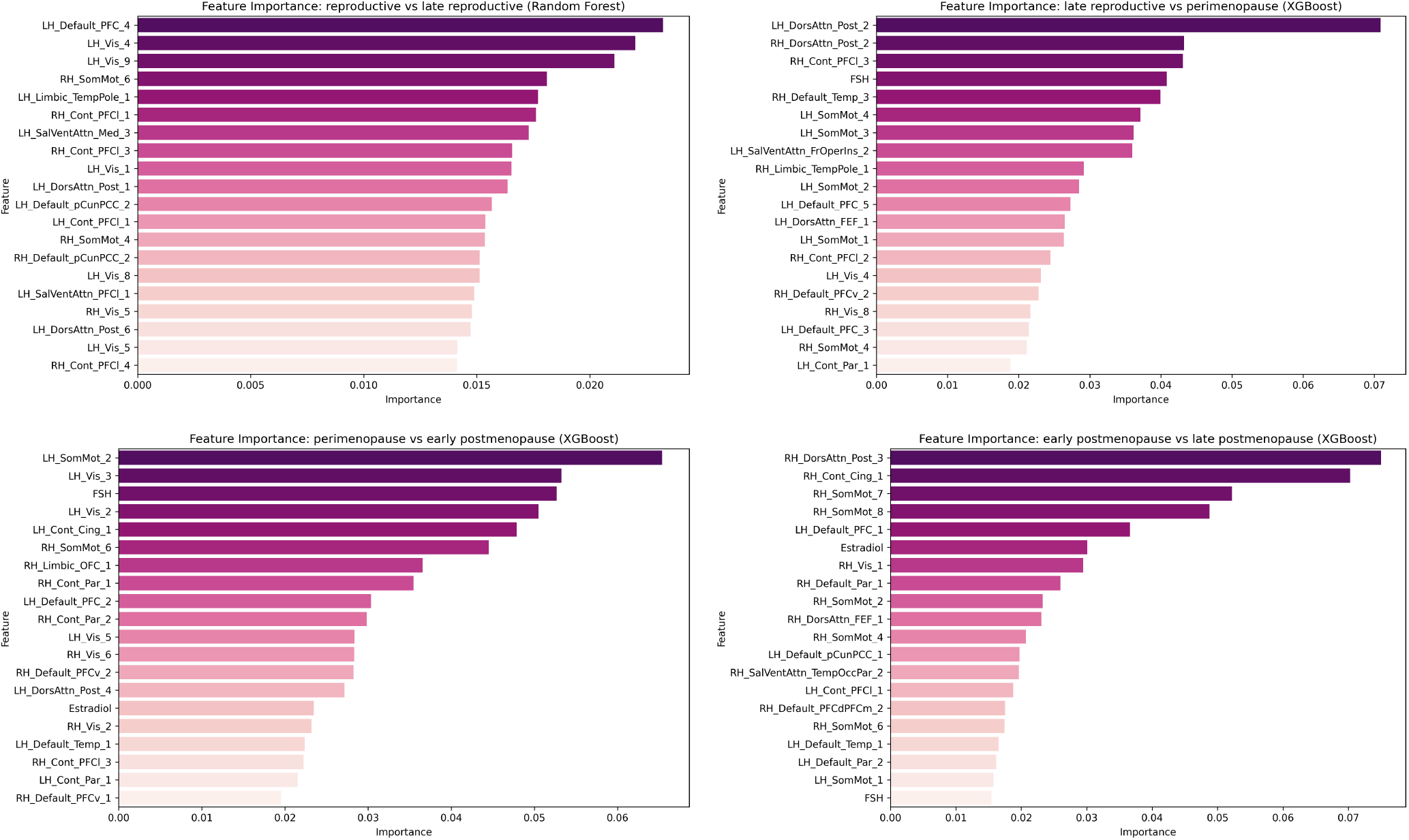
Top 20 most important features extracted from the hormone-brain models. The comparisons focus on transitions between consecutive stages, from reproductive to late postmenopause. This analysis shows the role of brain dynamic patterns and hormonal shifts in differentiating reproductive stages.

The Random Forest classifier best distinguished from reproductive to late reproductive stages. The top features included regions from the DMN, visual, somatomotor, salience, and frontoparietal control networks in both hemispheres and the left limbic temporal pole. However, no hormonal features were among the top predictors for this comparison.

XGB was the best-performing model for comparing late reproductive and perimenopause stages. The top predictive features included bilateral regions from the dorsal, frontoparietal control, DMN, and somatomotor networks. FSH levels were the fourth most important feature, indicating that rising FSH is a key marker of the transition to perimenopause.

For the comparison between perimenopause and early postmenopause, XGB was again the optimal model. The most predictive features were regions from the somatomotor, visual, frontoparietal control, DMN, and dorsal networks. FSH was the third most important feature, while estradiol ranked 15th. This suggests that both hormones are important for distinguishing perimenopause from early postmenopause.

XGB was superior in comparing early and late postmenopause stages. The top predictors included regions from the dorsal, frontoparietal control, somatomotor, and DMN networks. FSH was the 20th most important feature, while estradiol ranked 6th. This indicates that estradiol becomes a more salient marker than FSH for differentiating early from late postmenopause stages.

These results demonstrate the effectiveness of machine learning in identifying critical features of reproductive aging. While brain dynamic patterns are important across all comparisons, hormonal changes emerge as key predictors, especially the rise in FSH and decline in estradiol. These findings offer insights into the interplay between brain dynamics and hormonal fluctuations throughout women’s reproductive transitions.

## Discussion

To advance our understanding of women’s health, we explored brain dynamics evolution throughout reproductive stages. Our results revealed that whole-brain dynamics significantly change along adult women’s lifespan, from their reproductive years through the menopausal transition and beyond. Moreover, our analysis of resting-state networks showed significant reconfigurations across reproductive aging. Notably, results show that the most critical changes in brain dynamics occur from perimenopause to early postmenopause. Furthermore, machine learning classifiers trained with hormones and brain dynamics features demonstrated robust performance in distinguishing between reproductive stages.

The whole-brain analysis indicated that the late reproductive stage is characterized by the highest level of dynamical complexity across the functional network compared to other stages. This suggests that the human female brain experiences significant functional reorganization during this period, potentially reflecting the complex hormonal changes and physiological adaptations associated with this stage. Women in the reproductive stage showed higher complexity than early and late postmenopause stages but lower than late reproductive stage and perimenopause. These observations align with prior findings indicating increased dynamical complexity in early adulthood. For instance, neuroimaging investigations have consistently reported a zenith in network efficiency around midlife^10–12,30,31^. A study involving 60 reproductive-stage women aged between 18 and 35, scanned during three different phases of the menstrual cycle, corroborated this trend, highlighting increased complexity with age and when estradiol peaked^32^. Furthermore, Ballard et al.^20^ investigated differences in functional network segregation related to reproductive stages. They found a significant effect of reproductive stage vs. late postmenopausal across the whole network, suggesting that the menopausal transition is especially important for functional network communication.

Our analysis across resting-state networks indicates that women in the late reproductive stage generally exhibit the highest dynamical complexity, followed by those in perimenopausal and reproductive stages. This trend shows the significant influence of reproductive aging on brain network dynamics, with notable decreases in dynamical complexity observed in the early and late postmenopausal stages. Interestingly, perimenopausal women showed the highest dynamics in the frontoparietal control network compared to the other groups, although this finding was only marginally significant compared to the late reproductive stage. In particular, the complexity in this network increased from the reproductive stage to the late reproductive and then to the perimenopause stage, showing the peak during perimenopause, then significantly decreased in the early postmenopause and remained constant in the late postmenopause stage. However, in the salience network, brain dynamics decreased from the late reproductive stage to perimenopause and increased again from perimenopause to early postmenopause. The frontoparietal control and salience networks, along with the DMN, belong to the so-called triple network model^33,34^. The triple-network model posits that the salience network is crucial for processing relevant external events, which leads to the suppression of the DMN and modifies the temporal interactions among the DMN, salience, and frontoparietal control networks. These dynamic interactions result in the DMN disengaging from cognitive control systems, facilitating focused attention and working memory necessary for goal-directed behaviors. The model proposes that switching deficiencies in these three large-scale brain networks could play a significant role in psychopathology. While the DMN in our results showed the expected trend across reproductive stages, i.e., with the late reproductive stage showing the highest complexity levels compared to the other groups, the shifting disruptions between these networks during the menopausal transition could suggest an adaptative brain response to hormonal shifts during this critical period, which could reflect a compensatory mechanism during perimenopause before entering menopause.

Machine learning analysis based on brain dynamics and hormones demonstrated robust performance in classifying reproductive stages, achieving significant metrics. XGB demonstrated a slight advantage in more complex classification tasks (consecutive transitions), particularly distinguishing between late reproductive vs. perimenopause, late reproductive vs. early postmenopause, perimenopause vs. early postmenopause, and early postmenopause vs. late postmenopause stages. The high classification accuracies show the efficacy of using machine learning to assess reproductive aging. These results suggest that brain dynamics and hormone levels are promising biomarkers for reproductive stages, offering a novel approach to evaluating brain health in women. These results align with previous studies showing that perimenopausal and menopause women exhibit altered brain activity patterns compared to premenopausal women^4,16,18,19,21,35^. Moreover, studies utilizing RF and XGB have successfully classified menopausal women at high risk of depression^36^ and identified those with anxiety and depression with moderate accuracy^37^. These findings demonstrate the potential of machine learning in predicting mental health issues among menopausal women, suggesting that incorporating additional risk factors could enhance classification accuracy.

Several studies emphasized the importance of understanding the timing and trajectory of the menopausal transition for women’s health. The onset of menopause increases the risk of developing Alzheimer’s disease, likely due to the depletion of estrogen levels^38^. A study by Mosconi et al.^6^ found that the menopausal transition is associated with significant changes in Alzheimer’s disease biomarkers and cognitive performance, highlighting a female-specific risk for developing Alzheimer’s. Incorporating biomarkers related to neurodegeneration, such as tau and amyloid-*β* levels along with brain dynamics and hormone levels, into machine learning algorithms could help identify women at higher risk of developing Alzheimer’s disease.

The results of the most important features show the utility of machine learning for identifying key features that distinguish different stages of reproductive aging. We identified the most discriminative markers for each pairwise comparison across the transition from reproductive to late postmenopause. The top predictive features for classifying early and late reproductive stages included regions from the DMN, visual, somatomotor, salience, and frontoparietal control networks. However, no hormonal features were among the top predictors for this comparison. This suggests that brain dynamic patterns, particularly in the identified networks, are more important than hormonal changes for distinguishing between these groups. Moreover, classifiers were able to capture fluctuations in sex hormones (estrogen and FSH), which play a central role in the menopausal transition. The rise in FSH emerged as a key predictor for distinguishing late reproductive from perimenopause stages, as well as perimenopause from early postmenopause. This aligns with clinical evidence that increasing FSH levels are a hallmark of the menopausal transition^25^. Interestingly, estradiol became a more salient marker than FSH for differentiating early from late postmenopause stages. This likely reflects the continued decline in estradiol production by the ovaries as menopause progresses. Estradiol is known to have neuroprotective effects^39^, and its depletion during menopause has been linked to increased vulnerability to neuroinflammation and cognitive decline^40,41^. Our results suggest that estradiol levels may be particularly important for brain health in the later stages of menopause. In addition to hormonal changes, machine learning models identified key brain areas as predictors of the reproductive stage. Regions from the DMN, dorsal, somatomotor, and frontoparietal control networks emerged as important features across multiple comparisons. These networks are involved in diverse cognitive functions, including attention, sensorimotor integration, and higher-order cognition^17,34^. Alterations in the functional connectivity of these networks have been associated with cognitive changes during the menopausal transition. For example, the DMN has been linked to menopause-related memory complaints^42^, the dorsal network, involved in attentional processing, is altered in perimenopausal women, with enhanced functional connectivity, particularly in the right inferior parietal lobule and the right angular gyrus^17^

We want to acknowledge several limitations. The cross-sectional design limits the study of individual variability in each woman. Thus, longitudinal studies are required to understand the dynamic nature of reproductive aging and the interaction between hormonal fluctuations and brain dynamics. In addition, hormone levels were measured at a single time point, which may not fully capture their dynamic nature. This limitation is particularly notable for women aged 45 to 55, whose measurements were obtained only during the follicular phase, missing significant hormonal variations that occur throughout the menstrual cycle. Additionally, the analysis was limited to FSH and estradiol levels as hormonal features. Including other hormonal markers such as progesterone, luteinizing hormone (LH) and anti-Müllerian hormone (AMH) could lead to a more comprehensive understanding of hormonal dynamics in the context of reproductive aging. Future research could investigate the impact of other relevant variables that might influence whole-brain dynamic changes across the lifespan, such as demographic variables (e.g., educational level, employment status), number of pregnancies, ablation, polycystic ovary syndrome, cognitive indicators, etc.

In conclusion, this study shows that whole-brain dynamics and resting-state networks significantly reconfigurate throughout women’s lifespans. The results show the importance of considering hormonal and neuroimaging markers to study brain function across the female lifespan. This framework could be extended and used as a tool for personalized treatment by considering individual variability in hormone levels, brain dynamics, neurodegeneration biomarkers, and cognitive measures, particularly during the critical transition from perimenopause to menopause.

## Methods

### Participants

The HCP-A dataset comprises 725 healthy participants, including 406 women aged between 36 and over 100 years^26,43^. We specifically focused on resting-state fMRI data from women, excluding those who had undergone hysterectomy or ablation (61 participants), lacked a reproductive stage code (2), did not complete all four resting-state fMRI runs (2), or experienced irregular periods not due to natural menopause (10). As a result, our final sample consisted of 331 women. We obtained the reproductive stage from the data structure “mchq01” of the HCP-A dataset. Specifically, the HCP-A used the Menstrual Questionnaire, Menopause Screener, serum estradiol, FSH and STRAW+10 working group recommendations to define the reproductive stage. We categorized women into five groups based on their reproductive stage codes. Women with code 4.2 were classified as being in the reproductive stage, while those with code 4.1 were categorized as in the late reproductive stage. Women with codes 3, 2, 1.11, and 1.11004 were classified as perimenopausal. Those with codes 1.12, 1.12004, 1.13, and 1.13004 were categorized as early postmenopausal, whereas those with codes 1.14 and 1.14004 were categorized as late postmenopausal.

### MRI data acquisition and resting-state fMRI preprocessing

The HCP-A acquisition and fMRI preprocessing are described in full detail on the HCP website (https://www.humanconnectome.org/study/hcp-lifespan-aging). In brief, participants were scanned in a Siemens 3T Prisma with 80 mT/m gradients, a slew rate of 200T/m/s, and a 32-channel head coil. Neuroimaging data were acquired over two days in four sessions of 1-h. fMRI scans were acquired with a 2D-multiband (MB) gradient-recalled echo (GRE) echo-planar imaging (EPI) sequence (MB8, TR/TE = 800/37 ms, flip angle = 52°) and 2.0 mm isotropic voxels covering the whole brain (72 oblique-axial slices). Resting-state fMRI was acquired for 26 min in four runs of 6.5 min each, consistent with recent recommendations for obtaining robust connectivity estimates from fMRI data^44,45^. Participants looked at a small white fixation cross on a black background. They were instructed to stay still and awake while looking at the cross.

Resting-state fMRI data was preprocessed using the HCP pipeline, which employs FSL (FMRIB Software Library), FreeSurfer, and the Connectome Workbench software. This standardized preprocessing encompasses correction for spatial and gradient distortions, head motion correction, normalization, bias field removal, registration to the T1 structural image, transformation to the 2mm Montreal Neurological Institute (MNI) space, and FIX artifact removal^46^. Head motion parameters and artifacts were removed through ICA+FIX processing^47^ (Independent Component Analysis followed by FMRIB’s ICA-based X-noiseifier). fMRI data was mapped to CIFTI grayordinates space and functionally aligned across subjects, cleaned of spatially specific noise, concatenated, demeaned, and normalized across runs^48^.

### Brain parcellation

We employed custom MATLAB scripts using the ft_read_cifti function from the Fieldtrip toolbox^49^ to extract the time series of all grayordinates in each brain area in the Schaefer parcellation, according to the HCP CIFTI grayordinates standard space. The Schaefer cortical parcellation used^27^ divides the cortex into 100 brain areas clustered into 7 resting state networks: visual, somatomotor, dorsal attention, salience, limbic, frontoparietal control, and DMN. Details of the parcellation can be found at (https://github.com/ThomasYeoLab).

### Measure of dynamical complexity

We assessed the impact of intrinsic local perturbations, representing a brain area’s ability to propagate neural activity to other areas. This approach has been successfully used to distinguish different brain states in resting-state fMRI studies^11,12,22,32^. Specifically, node-metastability for each area is quantified as the standard deviation of global integration over time, capturing the level of whole-brain integration from spontaneous events. The algorithm identifies driving events in each region, which are transformed into a binary signal using a threshold^50^. Events are marked as 1 in the binary sequence if they exceed the threshold coming from below or are marked as 0 otherwise. When an area triggers an event, neural activity is assessed in all areas within a 4TR time window, and a binary matrix is constructed to represent connectivity. The global integration^51^ is then obtained from the largest sub-component to determine the network communication range for each event. This process is repeated for each spontaneous event to determine the node-metastability of brain areas over time. Higher values indicate increased dynamical complexity. Different levels of temporal diversity in a given brain area are related to its local functional variability or metastability, describing the versatility and dynamics of an area within the network. A complete description of the framework can be consulted in^12,22,32^.

### Statistical analyses

We employed Monte Carlo permutation tests to evaluate the differences in dynamical complexity across various reproductive stages within each network. For each pairwise comparison, we calculated the observed statistic as the absolute difference in means between the two groups. To establish a null distribution, we concatenated the data from the groups and performed 10,000 random shuffles, creating new null groups for each iteration. The null statistic was then computed as the absolute difference in means for these shuffled groups. The p-value for each comparison was determined by calculating the proportion of null statistics greater than or equal to the observed statistic. All reported p-values were adjusted for multiple comparisons using the False Discovery Rate (FDR) correction. This method was applied separately for each network when comparing between reproductive stages. Comparisons with FDR-corrected p-values less than 0.05 were considered statistically significant.

### Machine learning classifiers

We tested two approaches: 1) including 100 nodes for each woman (brain dynamics model), and 2) including FSH, estradiol and brain dynamics (brain-hormone model). We tested the group classification task using two distinct classifiers, *Random Forest*^28^ and *XGBoost*^29^. We employed a stratified 5-fold cross-validation strategy to ensure a suitable evaluation, preserving the distribution of the different reproductive stage groups across training and validation sets in each fold. We performed hyperparameter tuning for both classifiers using GridSearchCV to identify the optimal parameters, selecting the best based on the highest accuracy achieved during the grid search. For the Random Forest classifier, we tuned the following hyperparameters: n_estimators (50, 100, 200), max_depth (5, 10,, min_samples_split (2, 5, 10), and min_samples_leaf (1, 2, 4). For the XGBoost classifier, we tuned n_estimators (50, 100, 200), max_depth (3, 6, 10), and learning_rate (0.01, 0.1, 0.2). The choice of XGBoost hyperparameters was guided by recommendations in the XGBoost documentation and previous studies^29,52^. To address minority class imbalance, we employed the Synthetic Minority Over-sampling Technique (SMOTE)^53^. We evaluated the classifiers’ effectiveness using accuracy, precision, F1 score, and confusion matrix metrics. Missing hormone data, present for some women, were handled using mean imputation with the SimpleImputer strategy from scikit-learn^54^.

## Data and code availability

The HCP-A dataset is available at https://www.humanconnectome.org/study/hcp-lifespan-aging/article/lifespan-20-release-hcp-aging-hcp-development-data. The code supporting this work will be available on GitHub upon acceptance: https://github.com/aescrichs/reproductivestage-dynamicalcomplexity.

## Acknowledgements

A.E. was supported by the project eBRAIN-Health—Actionable Multilevel Health Data (ID 101058516), funded by the EU Horizon Europe, and by the Grant PID2022-136216NB-I00, funded by MICIU/AEI/10.13039/501100011033, and “ERDF A way of making Europe”, ERDF, EU. G.P. was supported by Grant PID2021-122136OB-C22 funded by MICIU/AEI/10.13039/501100011033 and by ERDF A way of making Europe. B.P. was supported by The European Research Council (ERC) Starting Grant 850953. P.R. acknowledges support by the Virtual Research Environment at the Charité Berlin – a node of EBRAINS Health Data Cloud, by EU Horizon Europe program Horizon EBRAINS2.0 (101147319), Virtual Brain Twin (101137289), EBRAINS-PREP 101079717, AISN – 101057655, EBRAIN-Health 101058516, Digital Europe TEF-Health 101100700, EU H2020 Virtual Brain Cloud 826421, Human Brain Project SGA2 785907; Human Brain Project SGA3 945539, German Research Foundation SFB 1436 (project ID 425899996); SFB 1315 (project ID 327654276); SFB 936 (project ID 178316478; SFB-TRR 295 (project ID 424778381); SPP Computational Connectomics RI 2073/6-1, RI 2073/10-2, RI 2073/9-1; DFG Clinical Research Group BECAUSE-Y 504745852, PHRASE Horizon EIC grant 101058240; Berlin Institute of Health amd Foundation Charité. Human Connectome Project in Aging (HCP-Aging) data used in this publication was supported by the National Institute On Aging of the National Institutes of Health under Award Number U01AG052564 and by funds provided by the McDonnell Center for Systems Neuroscience at Washington University in St. Louis. The HCP-Aging 2.0 Release data used in this report came from DOI: 10.15154/1520707.

## Author contributions statement

Conceptualization, methodology, and data curation: A.E. Data analysis: A.E, D.A., and G.P. Visualization: A.E, D.A., and G.P. Interpretation: All authors. Writing original draft: A.E. Writing, review, and editing: All authors.

## References

1. Hill, K. The demography of menopause. Maturitas 23, 113–127, DOI: 10.1016/0378-5122(95)00968-x (1996).

2. Brinton, R. D., Yao, J., Yin, F., Mack, W. J. & Cadenas, E. Perimenopause as a neurological transition state. Nat. Rev. Endocrinol. 11, 393–405, DOI: 10.1038/nrendo.2015.82 (2015).

3. Letenneur, L. et al. Education and the risk for Alzheimer’s disease: sex makes a difference. EURODEM pooled analyses. EURODEM Incidence Research Group. Am. J. Epidemiol. 151, 1064–1071, DOI: 10.1093/oxfordjournals.aje.a010149 (2000).

4. Kim, G.-W., Park, K., Kim, Y.-H. & Jeong, G.-W. Altered brain morphology and functional connectivity in postmenopausal women: automatic segmentation of whole-brain and thalamic subnuclei and resting-state fMRI. Aging 16, 4965–4979, DOI: 10.18632/aging.205662 (2024).

5. Nebel, R. A. et al. Understanding the impact of sex and gender in Alzheimer’s disease: A call to action. Alzheimer’s & Dementia 14, 1171–1183, DOI: 10.1016/j.jalz.2018.04.008 (2018).

6. Mosconi, L. et al. Increased Alzheimer’s risk during the menopause transition: A 3-year longitudinal brain imaging study. PloS One 13, e0207885, DOI: 10.1371/journal.pone.0207885 (2018).

7. Rosende-Roca, M. et al. The role of sex and gender in the selection of Alzheimer patients for clinical trial pre-screening. Alzheimer’s Res. & Ther. 13, 95, DOI: 10.1186/s13195-021-00833-4 (2021).

8. O’Neal, M. A. Women and the risk of Alzheimer’s disease. Front. Glob. Women’s Heal. 4, 1324522, DOI: 10.3389/fgwh.2023.1324522 (2023).

9. Gold, E. B. The Timing of the Age at Which Natural Menopause Occurs. Obstet. Gynecol. Clin. North Am. 38, 425–440, DOI: 10.1016/j.ogc.2011.05.002 (2011).

10. Damoiseaux, J. S. Effects of aging on functional and structural brain connectivity. NeuroImage 160, 32–40, DOI: 10.1016/j.neuroimage.2017.01.077 (2017).

11. Escrichs, A. et al. The effect of external stimulation on functional networks in the aging healthy human brain. Cereb. Cortex 33, 235–245, DOI: 10.1093/cercor/bhac064 (2023).

12. Escrichs, A. et al. Whole-Brain Dynamics in Aging: Disruptions in Functional Connectivity and the Role of the Rich Club. Cereb. Cortex 31, 2466–2481, DOI: 10.1093/cercor/bhaa367 (2021).

13. Shou, G., Yuan, H., Cha, Y.-H., Sweeney, J. A. & Ding, L. Age-related changes of whole-brain dynamics in spontaneous neuronal coactivations. Sci. Reports 12, 12140, DOI: 10.1038/s41598-022-16125-2 (2022).

14. Jacobs, E. G. Only 0.5% of neuroscience studies look at women’s health. Here’s how to change that. Nature 623, 667–667, DOI: 10.1038/d41586-023-03614-1 (2023).

15. Jett, S. et al. Ovarian steroid hormones: A long overlooked but critical contributor to brain aging and Alzheimer’s disease. Front. Aging Neurosci. 14, DOI: 10.3389/fnagi.2022.948219 (2022).

16. Liu, N., Zhang, Y., Liu, S., Zhang, X. & Liu, H. Brain functional changes in perimenopausal women: an amplitude of low-frequency fluctuation study. Menopause (New York, N.Y.) 28, 384–390, DOI: 10.1097/GME.0000000000001720 (2021).

17. Liu, N., Zhang, Y., Fu, W. & Liu, H. Functional changes in the dorsal attention network in perimenopausal women: a resting-state functional MRI study, DOI: 10.21203/rs.3.rs-4436654/v1 (2024).

18. He, L., Guo, W., Qiu, J., An, X. & Lu, W. Altered Spontaneous Brain Activity in Women During Menopause Transition and Its Association With Cognitive Function and Serum Estradiol Level. Front. Endocrinol. 12, DOI: 10.3389/fendo.2021.652512 (2021).

19. Zhang, Y., Fu, W. Q., Liu, N. N. & Liu, H. J. Alterations of regional homogeneity in perimenopause: a resting-state functional MRI study. Climacteric 25, 460–466, DOI: 10.1080/13697137.2021.2014808 (2022).

20. Ballard, H. K., Jackson, T. B., Symm, A. C., Hicks, T. H. & Bernard, J. A. Age-related differences in functional network segregation in the context of sex and reproductive stage. Hum. Brain Mapp. 44, 1949–1963, DOI: 10.1002/hbm.26184 (2022).

21. Zhang, S. et al. Aberrant Cerebral Activity in Early Postmenopausal Women: A Resting-State Functional Magnetic Resonance Imaging Study. Front. Cell. Neurosci. 12, DOI: 10.3389/fncel.2018.00454 (2018).

22. Deco, G. & Kringelbach, M. L. Hierarchy of Information Processing in the Brain: A Novel ‘Intrinsic Ignition’ Framework. Neuron 94, 961–968, DOI: 10.1016/j.neuron.2017.03.028 (2017).

23. Deco, G. & Kringelbach, M. L. Turbulent-like Dynamics in the Human Brain. Cell Reports 33, DOI: 10.1016/j.celrep.2020.108471 (2020).

24. Escrichs, A. et al. Unifying turbulent dynamics framework distinguishes different brain states. Commun. Biol. 5, 1–13, DOI: 10.1038/s42003-022-03576-6 (2022).

25. Harlow, S. D. et al. Executive Summary of Straw+10: Addressing the Unfinished Agenda of Staging Reproductive Aging. Climacteric : journal Int. Menopause Soc. 15, 105–114, DOI: 10.3109/13697137.2011.650656 (2012).

26. Bookheimer, S. Y. et al. The Lifespan Human Connectome Project in Aging: An overview. NeuroImage 185, 335–348, DOI: 10.1016/j.neuroimage.2018.10.009 (2019).

27. Schaefer, A. et al. Local-Global Parcellation of the Human Cerebral Cortex from Intrinsic Functional Connectivity MRI. Cereb. Cortex (New York, N.Y.: 1991) 28, 3095–3114, DOI: 10.1093/cercor/bhx179 (2018).

28. Breiman, L. Random Forests. Mach. Learn. 45, 5–32, DOI: 10.1023/A:1010933404324 (2001).

29. Chen, T. & Guestrin, C. XGBoost: A Scalable Tree Boosting System. In Proceedings of the 22nd ACM SIGKDD International Conference on Knowledge Discovery and Data Mining, 785–794, DOI: 10.1145/2939672.2939785 (2016).

30. Cao, M. et al. Topological organization of the human brain functional connectome across the lifespan. Dev. Cogn. Neurosci. 7, 76–93, DOI: 10.1016/j.dcn.2013.11.004 (2014).

31. Zhao, T. et al. Age-related changes in the topological organization of the white matter structural connectome across the human lifespan. Hum. Brain Mapp. 36, 3777–3792, DOI: 10.1002/hbm.22877 (2015).

32. Avila-Varela, D. S. et al. Whole-brain dynamics across the menstrual cycle: the role of hormonal fluctuations and age in healthy women. npj Women’s Heal. 2, 1–9, DOI: 10.1038/s44294-024-00012-4 (2024).

33. Menon, V. Large-scale brain networks and psychopathology: a unifying triple network model. Trends Cogn. Sci. 15, 483–506, DOI: 10.1016/j.tics.2011.08.003 (2011).

34. Menon, V. 20 years of the default mode network: A review and synthesis. Neuron 111, 2469–2487, DOI: 10.1016/j.neuron.2023.04.023 (2023).

35. Berent-Spillson, A. et al. Hormonal Environment Affects Cognition Independent of Age during the Menopause Transition. The J. Clin. Endocrinol. Metab. 97, E1686–E1694, DOI: 10.1210/jc.2012-1365 (2012).

36. Ali, M. M. et al. Development and performance analysis of machine learning methods for predicting depression among menopausal women. Healthc. Anal. 3, 100202, DOI: 10.1016/j.health.2023.100202 (2023).

37. Martin, C. M., Larroy, C., López-Picado, A. & Fernández-Arias, I. Accuracy of the Menopause Rating Scale and the Menopause Quality of Life Questionnaire to discriminate menopausal women with anxiety and depression. Menopause 26, 856, DOI: 10.1097/GME.0000000000001338 (2019).

38. Christensen, A. & Pike, C. J. Menopause, obesity and inflammation: interactive risk factors for Alzheimer’s disease. Front. Aging Neurosci. 7, DOI: 10.3389/fnagi.2015.00130 (2015).

39. Wise, P. M., Dubal, D. B., Wilson, M. E., Rau, S. W. & Böttner, M. Minireview: Neuroprotective Effects of Estrogen—New Insights into Mechanisms of Action. Endocrinology 142, 969–973, DOI: 10.1210/endo.142.3.8033 (2001).

40. Guerrero-González, C., Cueto-Ureña, C., Cantón-Habas, V., Ramírez-Expósito, M. J. & Martínez-Martos, J. M. Healthy Aging in Menopause: Prevention of Cognitive Decline, Depression and Dementia through Physical Exercise. Physiologia 4, 115–138, DOI: 10.3390/physiologia4010007 (2024).

41. Nerattini, M. et al. Systematic review and meta-analysis of the effects of menopause hormone therapy on risk of Alzheimer’s disease and dementia. Front. Aging Neurosci. 15, DOI: 10.3389/fnagi.2023.1260427 (2023).

42. Thurston, R. C., Maki, P. M., Derby, C. A., Sejdić, E. & Aizenstein, H. J. Menopausal Hot Flashes and the Default Mode Network. Fertility sterility 103, 1572–1578.e1, DOI: 10.1016/j.fertnstert.2015.03.008 (2015).

43. Harms, M. P. et al. Extending the Human Connectome Project across ages: Imaging protocols for the Lifespan Development and Aging projects. NeuroImage 183, 972–984, DOI: 10.1016/j.neuroimage.2018.09.060 (2018).

44. Laumann, T. O. et al. Functional System and Areal Organization of a Highly Sampled Individual Human Brain. Neuron 87, 657–670, DOI: 10.1016/j.neuron.2015.06.037 (2015).

45. Glasser, M. F. et al. The Human Connectome Project’s neuroimaging approach. Nat. Neurosci. 19, 1175–1187, DOI: 10.1038/nn.4361 (2016).

46. Smith, S. M. et al. Resting-state fMRI in the Human Connectome Project. NeuroImage 80, 144–168, DOI: 10.1016/j.neuroimage.2013.05.039 (2013).

47. Salimi-Khorshidi, G. et al. Automatic denoising of functional MRI data: Combining independent component analysis and hierarchical fusion of classifiers. NeuroImage 90, 449–468, DOI: 10.1016/j.neuroimage.2013.11.046 (2014).

48. Glasser, M. F. et al. The minimal preprocessing pipelines for the Human Connectome Project. NeuroImage 80, 105–124, DOI: 10.1016/j.neuroimage.2013.04.127 (2013).

49. Oostenveld, R., Fries, P., Maris, E. & Schoffelen, J.-M. FieldTrip: Open source software for advanced analysis of MEG, EEG, and invasive electrophysiological data. Comput. Intell. Neurosci. 2011, 156869, DOI: 10.1155/2011/156869 (2011).

50. Tagliazucchi, E., Balenzuela, P., Fraiman, D. & Chialvo, D. R. Criticality in Large-Scale Brain fMRI Dynamics Unveiled by a Novel Point Process Analysis. Front. Physiol. 3, DOI: 10.3389/fphys.2012.00015 (2012).

51. Deco, G., Tononi, G., Boly, M. & Kringelbach, M. L. Rethinking segregation and integration: contributions of whole-brain modelling. Nat. Rev. Neurosci. 16, 430–439, DOI: 10.1038/nrn3963 (2015).

52. Friedman, J. H. Greedy function approximation: A gradient boosting machine. The Annals Stat. 29, 1189–1232, DOI: 10.1214/aos/1013203451 (2001).

53. Chawla, N. V., Bowyer, K. W., Hall, L. O. & Kegelmeyer, W. P. SMOTE: Synthetic Minority Over-sampling Technique. J. Artif. Intell. Res. 16, 321–357, DOI: 10.1613/jair.953 (2002).

54. Pedregosa, F. et al. Scikit-learn: Machine learning in python. J. machine Learn. research 12, 2825–2830 (2011).

